# Physicochemical Characterization of Stingless Bees’ (*Meliponula beccarii L)* Honey from Wonchi District, Southwest Shewa Zone, Ethiopia

**DOI:** 10.64898/2026.04.01.715950

**Authors:** Samuel Adugna Gedefa, Debebe Landina Lata

## Abstract

This study was aimed at characterizing the physicochemical analysis of stingless bees’ honey (SBH) in the Wonchi district, Southwest Shewa Zone, Ethiopia. In this study, a total of 30 stingless bees’ honey samples were collected from Damu Dagele, Fite Wato, and Warabu Messe sites from underground soils through an excavation of natural nests. Physicochemical characterization of properties and proximate analysis of the honey were performed. The result showed a total mean of 20.12±1.14% moisture content, 8.62±2.73 meq./kg free acidity, 1.8±0.52 mS/cm electrical conductivity, 3.39±0.32 pH, 40.52±6.61 mg/kg HMF, 0.83±0.33% ash, 0.56±0.25% protein, 0.56±0.24% fat, and 0.59±0.23% WISC for physicochemical properties of stingless bees’ honey. Among sugar profiles of SBH, fructose constituted the highest proportion at 18.87 g per 100 g (53.87%), while sucrose exhibited the lowest concentration at 5 g per 100 g (14.33%). The result showed that the highest constituted mean of mineral composition was observed with potassium (K) of 16.64±0.257 mg/kg, while magnesium (Mg) showed the lowest concentration of 3.48±0.17 mg/kg. A substantial correlation was observed between K and Mg, with a correlation coefficient of 0.72 and 0.72, and similarly between K and Calcium (Ca); the correlation was highly significant, exhibiting a correlation coefficient of 0.65. Furthermore, the correlation between fatty and other physicochemical and proximate analyses showed very insignificant correlations. In general, this study showed that the SBH produced in the current study area has good physicochemical properties and moisture and contains high-quality honey, which may help its traditional medicinal uses. The findings of the study further suggests the potentiality of the area for quality honey, and to easily locate priority areas for stingless bee conservation, further detailed studies of other stingless species’ honey medicinal values are recommended.

## 1. INTRODUCTION

Stingless bee honey (SBH) is a type of honey bee that has no sting attribute, as its name suggests, and is a member of the same Hymenoptera family (Rasmussen & Cameron, 2010). Stingless bees (SB) are small, all-black insects found in tropical and subtropical areas of the world. They are unique in generating three different types of products: honey, propolis, and bee bread (Nurhadi, 2016). Stingless bee honey is characterized by having different types of physicochemical and proximate composition (Mathiasson, 2015). Honey is a natural sweetener with roughly 200 different chemical compositions, including 80-85% carbohydrates like fructose and glucose (Rao *et al*., 2016). They also contain water, proteins, and amino acids, as well as ash, enzymes, vitamins, and phenolic compounds, which make up nearly 15–20% of honey (Rao *et al*., 2016). The composition of stingless bee honey differs from other species and is more valuable (Souza *et al.,* 2006). Its major composition includes simple sugars (fructose and glucose), organic acids, phenolic compounds, proteins, amino acids, and minerals (Ismail, 2016). Furthermore, stingless bee honey contains polyphenols, vitamins, enzymes, amino acids, and minerals (Kahraman *et al*., 2010). However, the composition of stingless bee honey varies according to the floral source and origin (Lusby *et al*., 2002). When compared to honey from honey bee (*Apis mellifera),* stingless bee honey has higher acidity levels, reducing sugars, distinct sweetness, flavor, texture, and scent (Ávila *et al*., 2018; Souza *et al*., 2006). Approximately 200 different chemical compositions found in SBH are widely utilized for a variety of purposes (Rao *et al*., 2016). However, the composition of honey varies based on the kinds of plants and the nectar that the bee eats.

The most common SBH quality standard is moisture content, which is followed by free acidity, sugar profile, pH, hydroxymethyl furfural (HMF), ash content, and electrical conductivity (Chuttong *et al.,* 2016). The amount of moisture in honey is crucial because it could affect several other factors, including the amount of sugar, HMF, and microbiological characteristics. According to Chuttong *et al. (*2016), SBH is believed to have more moisture, more electrical conductivity, less enzyme activity, more free acidity, and less glucose and fructose than *Apis mellifera* honey. It also contains carbohydrates, proteins, amino acids, lipids, minerals, vitamins, fragrance compounds, flavonoids, organic acids, pigments, enzymes, and phytochemicals. The content and properties of honey are influenced by climate and environmental conditions (Iglesias *et al*., 2012). SBH’s color varies from colony to colony, and its flavor is influenced by the plants they collect nectar from (Kulkarni *et al.,* 2017). The physicochemical and proximate composition of honey is also responsible for its antimicrobial activities.

SBH products, such as honey and cerumen, have been used as a source of income for generations. Stingless bees play an important role in the economy, ecology, and culture. They act as the main pollinators for many wild and cultivated plants in the tropics (Slaa *et al.,* 2006). They could act as pollinators of a wide range of plants, including vegetables, pulses, oilseeds, fruit trees, and plantations throughout the tropical and subtropical parts of the world (Heard, 1999). In Ethiopia, stingless bee honey is available in a limited area. SBH in Ethiopia are not well known and probably get attention due to their limited distribution in the country. Thus, the present study aimed to assess the microbiological and physicochemical qualities of SBH’s collected from Wonchi, one of the producing areas of the country.

Honey production and stingless honey bees are environmentally friendly practices and relatively easy to engage in. Despite its high demand and medicinal value, the issue of its quality and authenticity remains an important factor in its consumption and marketing due to the scant knowledge about its production system and composition. As a result, the proximate composition property of SBH is not yet characterized and documented, even to set its quality standard both for nutritional and medicinal value. Eventually, the result helps set SBH quality standards and its characterization, particularly in the identification of Ethiopian stingless bee honey. Therefore, the current study aims to evaluate the proper physicochemical characteristics of honey from native stingless bees (*Meliponula beccarii*) of Ethiopia and determine some of its quality levels in comparison with *A. mellifera* honey. Although stingless bee honey has wide advantages, it is rare in Ethiopia and available only in a limited area. Furthermore, SBH is declining throughout the country, and there is a lack of scientific research in these areas. Hence, the present study aimed to assess the physicochemical properties and proximate analysis of stingless bees’ honey from the southwest Shewa Zone, Ethiopia.

## 2. MATERIAL AND METHODS

### 2.1. Description of the Study Sites

The study was carried out in Wonchi district, which is located in South West Shewa Zone, Oromia Regional State, between October and January 2024. It is located about 37km West of Woliso, the capital of the South West Shewa Zone administrative, and 27 km southeast of Ambo, the capital town of West Shewa (**Figure. 1**). The district is found 150 km west of Addis Ababa, the capital city of Ethiopia. The geographic location of the district lies between 08°41′N and 037°53′E, altitudinal variation of the district extends from 1700 to 3387 m a.s.l. The total area of the Woreda is 475.6 km2. The total human population of the district is about 119,736. The district is bordered by seven other districts: Toke Kutaye (North West), Ambo and Dendi (North), Dawo (North East), Woliso (East), Amaya (South West), and Goro (South). The district is typically characterized by Wonchi Lake, which is a source of ecotourism for the country and on which the government currently has a big project nationally (Dida *et al.,* 2020).

**Figure 1:**
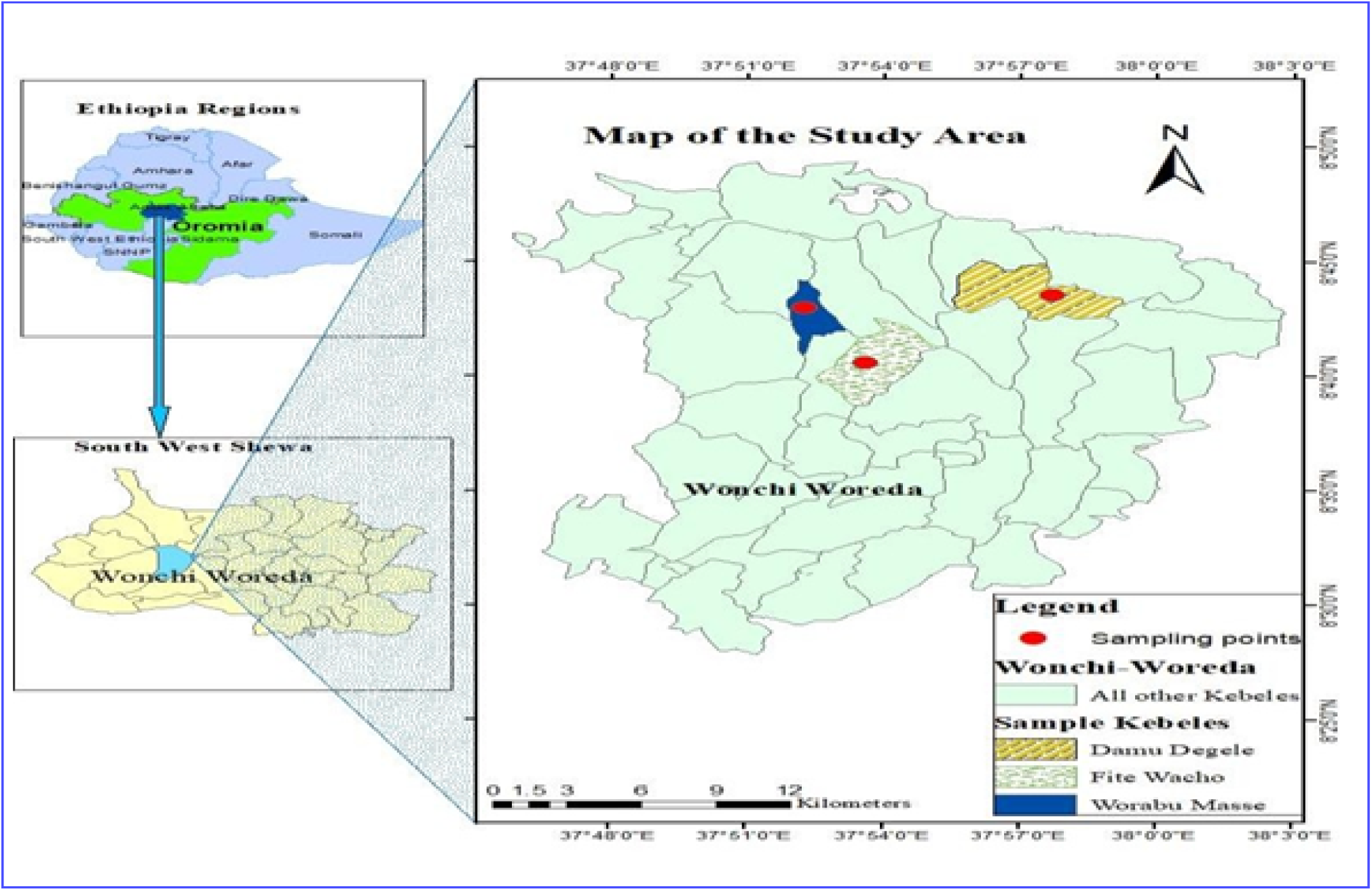
Map of the study area, Wonchi district

### 2.2 Climate of the Study Area

The annual rainfall of the Wonchi district reached up to 1142 mm in some peak years. The average monthly high and low temperatures of the area are 22.7°C and 11.9°C, respectively. The average annual temperature of the district was 16.9°C, which slightly varies from year to year. The rainfall pattern shows low rainfall in December and February, which gradually increases to the peak period in July and then decreases in November and December. The district is known predominantly for growing barley and wheat, multipurpose trees, herbs, shrubs, the production of honey, and traditional beverages. Moreover, the district’s plant diversity was very important for the production of honey and stingless honey. Particularly, SBH is found in a few areas of Ethiopia and is rarely found in other countries (Meragiaw *et al*., 2022).

### 2.3 Sample Collection

A total of 30 stingless bee honey samples of 300ml honey were collected from Warabu masi, Damu Dagale, and Fitte wato (10 samples from each kebele). These three sites were selected based on a survey conducted and information obtained from the local informant about the presence of SBH. The SBH samples were collected aseptically by using a sterile bottle placed in an icebox and transported to Wolkite University Biology and Biotechnology Departments Laboratories for analysis of microbiological quality, physicochemical properties, proximate composition, and antimicrobial activities. The samples were kept at 4°C until the activities were conducted.

### 2.4 Physicochemical and Proximate Composition Analysis

#### 2.4.1 pH and Free Acidity Determination

The pH and free acidity were determined according to Tri’s (2009) technique. Briefly, for pH determination, 10 g of stingless bee honey and 75 ml of distilled water were used to dissolve the SBH sample and then dipped in a pH meter individually. The free acidity of the SBH sample was determined by dissolving 10g of SBH in 75 mL of distilled water. Then, titrate with 0.1M NaOH solution (pH = 8.3) after adding 3 to 5 drops of 1g/100ml phenolphthalein indicator. Then, after, free acidity is calculated by: Free Acidity=10 × Volume of 0.1M NaOH used (Sadler & Murphy, 2010).

#### 2.4.2. Determination of Moisture and Total Solid Content

The moisture content of SBH would be determined by measuring an empty beaker (W1) that has been dried in a 105 °C oven for two hours. Then, 10 g of SBH samples were weighed separately and added to pre-weighed dry beakers (W2) before being dried in an oven at 105 °C for 2 h. The samples were moved to a desiccator after 2 hr of drying, and the weight (W3) was calculated by the Association of Official Analytical Chemists (Nielsen & Bradley, 2010).

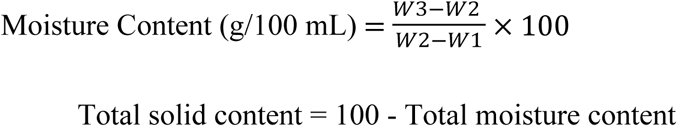

#### 2.4.3 Determination of Water-insoluble and Soluble Solid Contents

The SBH samples’ water-insoluble and soluble solid contents were determined using Ethiopia’s Quality and Standards Authority’s protocols (Zhang *et al*., 2022). Briefly, 20 g of SBH was dissolved in 200 mL of boiling water at 80°C. To remove sugar from the sample solution, it was filtered through a previously dry, weighed fine sintered glass crucible, and washed extensively with hot water (80 °C) 3-5 times. The crucible was then dried at 130 °C for 1h, cooled, and the weights calculated as follows:

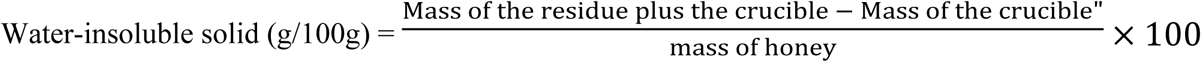

Water-soluble solid = Total solid content - Water-insoluble solid.

#### 2.4.4 Determination of Electrical Conductivity (EC)

The EC would be measured by dissolving 20g of SBH in 100 mL of distilled water, and the result would be recorded after inserting a digital multi-parameter into the mixture (Zappalà *et al*., 2005).

#### 2.4.5 Determination of Ash content

The ash content was determined using the Association of Official Analytical Chemists method. Accordingly, 5 g of SBH sample was weighed in a silica crucible and ignited in a muffle furnace for about 3 h at 550°C. The following formula would calculate ash content (Noh & Schwarz, 1990)

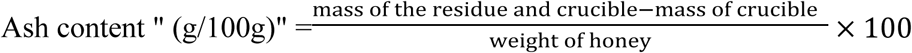

#### 2.4.6 Determination of Hydroxy-Methyl-Furfural (HMF) Content

The HMF content was determined based on the procedure of (Makawi *et al*., 2009). Accordingly, 5 g SBH was dissolved in 25 ml distilled water and treated with 0.5ml of Carrez I (Dissolve 15g potassium Ferro cyanide (K^4^F^e^(CN)_6_. 3H_2_O) in distilled water and dilute to 100 mL) and 0.5mL Carrez II solution (Dissolve 30g zinc acetate (Zn (CH_3_CO_2_)_2_·2H_2_O) in distilled water and dilute to 100ml) and dilute to a volume of 50ml with distilled water. The homogenate was filtered through 26 x 31mm filter paper, discarding the first 10ml filtrate. Then, a 5mL filtrate was piped into each of the two test tubes. To the first test tube (sample), 5mL distilled water was added, and 5ml NaHSO_3_ solution (Dissolve 0.2% NaHSO_3_ and dilute to 100ml water) to other (reference) and mixed using a vortex mixer, and the absorbance of the sample was determined against reference at 284 and 336 nm. Eventually, HMF was calculated using the following formula: (Makawi *et al*., 2009)

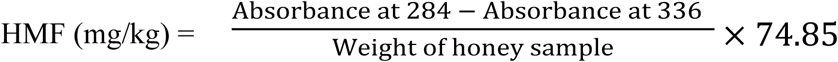

#### 2.4.7 Determination of Total Protein Content

The Kjeldahl approach was used to calculate total protein contents following the procedure of the Association of Official Analytical Chemists (1990). Briefly, 2 g of the SBH sample, a catalyst amount of 0.2g of CuSO_4_, and 1g of K_2_SO_4_ were separately added in a micro-Kjeldahl flask. Then, 15 mL of concentrated H_2_SO_4_ (w/v) was further added to each sample. These samples were digested at 420 °C for 75 min. The digested samples were then diluted with 50 mL of distilled water (v/v), and the micro-Kjeldahl flask was attached to the distillation unit. Then, 45 ml of 15 mol/L NaOH was added, and the sample distillation released ammonia, which was collected into the boric acid solution. (In relation to the amount of nitrogen) Borate anion) was titrated with standardized 0.1mol/L H_2_SO_4_. An automatic Kjeldahl analyzer (Velp ScientificaTM UDK, F30200150) method of determination is used to determine the percent protein content (Campos *et al*., 2008).

#### 2.4.8 Determination of Total Fat Content

The total fat content was determined according to the procedure of the Association of Official Analytical Chemists (1990). Briefly, 5 g of the SBH sample was taken and mixed thoroughly with 2 ml of 99% ethyl alcohol (w/v) in the extraction apparatus. Then, diluted with 10ml of HCl (44%, v/v) and mixed well. The hydrolyzed fat was extracted with petroleum ether (100 mL) for a minimum period of 4h in the soxhlet extractor. The petroleum ether was evaporated from the extract, and the fat was dried to a constant weight at 100 °C for 90 min (Messner *et al*., 2014). The total fat was calculated as follows:

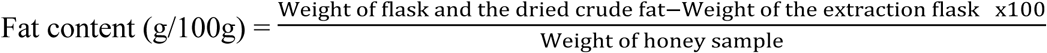

#### 2.4.9 Determination of Sugar Contents

Sugar analysis was conducted based on the international SBH commission Nordin *et al*. (2018) methods. Accordingly, each sugar standard solution of fructose (2 g), glucose (1.5 g), and sucrose (0.25 g) was dissolved in 20 mL of HPLC-grade water. A total of 25 mL methanol was transferred into a 100 mL calibrated flask. Then, a standard solution was poured into a flask-containing methanol and filled to the mark with HPLC-grade water. This solution be filtered through a 0.45-μm membrane filter syringe, and the filtrate will be poured into a sample vial (Agilent, 1260 infinity, USA) for injection. HPLC conditions flow rate: 1.5 mL/min; mobile phase: acetonitrile: water (75:25, v/v); column and detector temperature: 30 °C; and sample volume: 10 μl was used for HPLC separation, and peaks be identified based on their retention times. The SBH samples was prepared by dissolving 5 g in 40 mL of HPLC-grade water. A total of 25 mL of methanol was pipetted into a 100 mL volumetric flask, transferred SBH solution quantitatively to the flask, and then filled to the mark with HPLC-grade water. After that, the SBH solution was filtered through a membrane filter syringe and collected in the sample vial *(Rossmann et al*., 1997).

#### 2.4.10 Determination of Mineral Content

Minerals’ standards were prepared at different concentrations in a 1% HNO_3_ (v/v) medium. The mineral contents of SBH were determined by Agilent microwave plasma atomic emission spectroscopy (MP-AES). Accordingly, 0.5 g of each SBH sample was digested with 4 mL of HNO_3_ (65% v/v) and 1 mL of H_2_O^2^ (30 % v/v). All samples were microwave digested using the Speed wave four microwave system (Agilent 4200 MP-AES, USA) in vessels (40 mL, 230 °C, and 40 bar of pressure), according to the following heating program: (i) heating ramp: 120 °C, 5 min, (ii) heating plateau: 120 °C, 5 min, (iii) heating ramp: 160 °C, 5 min, (iv) heating plateau: 160 °C, 5 min, (v) heating ramp: 200 °C, 5 min, (vi) heating plateau: 200 °C, 5 min, (vii) cooling step: 3 min at 22 °C and diluted with 50 mL deionized ultrapure water. Finally, this solution was used for elemental analysis (Masamba & Kazombo, 2010).

### 2.5 Data Analysis

In order to analyze the data, SPSS version 20.0 was used. The significance difference in the experimental treatment was analyzed using one way analysis of variance (ANOVA). P ≤ 0.05 was considered statistically significant. The correlation between physicochemical, proximate, and microbes was done using Person correlations.

## 3. RESULTSAND DISCUSSION

### 3.1. Physicochemical Properties and Proximate Composition Analysis

The current study identified different composites from SBH. Accordingly, moisture, fat, protein, magnesium, potassium, calcium, and ash were identified from stingless bees’ honey collected from three different kebeles of Wonchi Woreda (**Table 1**). Altogether, the composites for stingless bee honey collected from Damu Degele, Fite Wacho and Worabu Messe varied in the following ranges: moisture (17.82-21.54%), fat (0.27-0.96%), protein (0.44-0.87%), Mg (2.46-4.37%), K (14.5-18.34%), Ca (4.16-7.40%), and ash (0.24-1.14%). From these composites, moisture content, fat, protein, K, and Ca are significantly higher in stingless bees’ honey of Damu Degele and Worabu Messe than in Fite Wacho (P=0.00, P=0.04, P=0.01 and P= 0.00, P=0.00, respectively). In addition, magnesium is significantly higher in stingless bees’ honey collected from Damu Degele (p=0.00). Such variability in chemical composition among honey collected from different locations is expected since it is influenced by different conditions, including geographical location, source of nectar, seasonal and environmental factors, and handling techniques ( Silva *et al.,* 2013). However, there was no significant difference among the ash component of stingless bees’ honey collected from the three locations (*p=0.12*). However, the overall mean value of moisture content of SBH (20.12%) of the recent study was lower than the previously reported moisture content against SBH produced in Ethiopia (25.10-35%) (Gela *et al.,* 2021), Venezuela (26-42%), and Brazil (25-36%). In contrast to this study, the mean value of moisture content against the SBH (20.12%) is similar to the international standard moisture content for *A. mellifera* honey, which permits a maximum moisture content of 20% (Nordin *et al.,* 2018).

**Table 1:** Physicochemical and proximate analysis of SBH.

| Parameter | Damu Degele | Fite | Worabu | Total | LSD | CV% | P-value |
| --- | --- | --- | --- | --- | --- | --- | --- |
|  |  | walo | Messe |  |  |  |  |

|  | Mean ± SD | Mean ± SD | Mean ± SD | Mean±SD |  |  |  |
| --- | --- | --- | --- | --- | --- | --- | --- |
| Fat (%) | 0.47<sup>a</sup>±0.20 | 0.27<sup>b</sup>±0.24 | 0.96<sup>a</sup>±0.3 | 0.56±0.24 | 0.54 | 47.49 | 0.04 |
| Protein (%) | 0.70<sup>a</sup>± 0.21 | 0.44<sup>b</sup>±0.24 | 0.87<sup>a</sup>±0.3 | 0.56±0.25 | 0.24 | 18.49 | 0.01 |
| Ash(mS/cm) | 1.14<sup>a</sup>± 0.63 | 0.24<sup>a</sup>±0.15 | 1.11<sup>a</sup>±0.2 | 0.83±0.33 | 1.03 | 61.81 | 0.12 |
| Moisture (g/100g) | 21.54<sup>a</sup>± 1.92 | 17.82<sup>b</sup>±0.5 | 20.99<sup>a</sup>±1.0 | 20.12±1.14 | 1.87 | 4.67 | 0 |
| PH | 3.68<sup>a</sup>± 0.34 | 2.95<sup>b</sup>±0.23 | 3.54<sup>a</sup>±0.4 | 3.39±0.32 | 0.25 | 3.65 | 0 |
| HMF | 44.85<sup>a</sup>± 7.12 | 32.55<sup>b</sup>±6.7 | 44.16<sup>a</sup>±6.0 | 40.52±6.61 | 8.89 | 10.99 | 0.02 |
| EC | 2.15<sup>a</sup>± 0.39 | 1.30<sup>b</sup>±0.38 | 1.95<sup>a</sup>±0.8 | 1.8±0.52 | 0.25 | 7.01 | 0 |
| Free acidity (meq/kg) | 9.18<sup>a</sup>± 2.45 | 5.92<sup>b</sup>±2.37 | 10.76<sup>a</sup>±2. | 8.62±2.73 | 2.81 | 16.35 | 0.02 |
| WISC (%) | 0.87<sup>a</sup>± 0.29 | 0.24<sup>b</sup>±0.2 | 0.67<sup>a</sup>±0.2 | 0.59±0.23 | 0.25 | 21.46 | 0 |

*EC=Electric conductivity; HMF=Hydroxyl Methyl Furfural; WISC=Water-Insoluble Solids Content;* *CV=Coefficient of variation, SD=Standard Deviation, LSD=Least Significant Difference; and superscripts in* *the column followed by the different letter are significantly different at the 5% level by Fisher's LSD.*

This study showed that the ash content in SBH varied significantly, ranging from 0.24 to 1.14%. This result is relatively higher when compared to previous studies done in Ethiopia (0.21-0.56%) Gela *et al. (*2021), Malaysia (0.04-0.19%) Hasali *et al*. (2018), and Brazil (0.19-0.33%) (Rosenthal *et al*., 2016). Such variations among the study locations and other previous studies could be due to variations in botanical origin, diversities of SHB, sampling location, and processing & handling of SBH (Biluca *et al*., 2016). In addition, the average ash content (0.83%) of SHB in this study is relatively higher than *Apis* honey (0.14-0.3%). This result suggests that honey produced from native SHB in Ethiopia is richer in mineral contents and, therefore, could be good nutritional compositions and medicinal value. This high mineral content might be associated with the origin of SBH that is harvested from the ground, where the mineral content is expected to be higher than in the beehives.

Protein (0.44-0.87%) and fat (0.27-0.96%) contents of SBH in this study expressly varied among the study sites. Such differences in protein content could be attributed to the type of flora and types of honey from various botanical origins (Saxena *et al.,* 2010). In addition, protein in the honey mainly consists of enzymes and free amino acids that can easily be affected during processing and climate change, which contributes to the variations of the protein content in the honey samples (Access *et al.,* 2014). However, the protein content of the present study against SBH is almost similar to the study done by Hasali *et al*. (2018), which ranges from 0.34 to 0.69%. On the other hand, the considerable fat content in this study is in contrast to other studies done by Hasali *et al*. (2018) in which the honey harbored zero fat content. Changes in temperature, precipitation patterns, and overall environmental conditions can affect the availability of floral resources for honeybees, which in turn can influence the protein content of the honey they produce.

Concerning the physicochemical properties of SBH, six physicochemical components, namely TSS, pH, HMF, EC, free acidity, and WISC, were analyzed. Similar results were reported for all three sampled kebeles. All those six physicochemical characteristics of the SBH were significantly lower in Fite Wacho when compared to the other two locations (p=0.00, p=0.00, p=0.02, p=0.00, p=0.02, and p=0.00, respectively). However, there was no significant difference between the physicochemical properties of the SBH collected from Damu Degele and Worabu Messe. Overall physicochemical parameters for stingless bee honey collected from the three locations range from 57.51 to 70.37 for TWSS, 3.33 to 4.88 for pH, 32.55 to 44.85 mg/kg for HMF, 1.30 to 2.15 mS/cm for EC, 5.92 to 10.76 meq/kg for free acidity, and 0.24 to 0.87% for WISC.

Low pH enhances SBH’s shelf life and can also inhibit the presence and growth of microorganisms. Acidity, which is a quality parameter that indicates the presence of organic acids in honey, varied among the three kebeles in this study. This variation might be attributed to the floral origin, location, and its management as reported in the previous study (Nordin *et al.,* 2018).

The recent acidity value (8.62 meq/kg) for the SBH was lower when compared to the previous study by Gela *et al*. (2021) with a 17.3 meq/kg value. However, this average value of acidity for the SBH is laid within the acceptable limit of international standard values (<50 meq/kg) for *Apis* honey. The mean average electrical conductivity (which is related to the number of organic acids, protein, and mineral salts in honey) for the present study is far greater than the international standard value (<0.8 mScm⁻¹), the overall average value of SBH collected from Ethiopia (0.21 mScm⁻¹), and Brazil (0.74 mScm⁻¹) (Gela *et al.,* 2021).

Another physicochemical property that was characterized in this study is hydroxymethyl furfural (HMF), which is made from the chemical reaction of some acids and sugars, and it is used as an indicator of honey freshness and good quality (Nordin *et al.,* 2018). A considerable amount of HMF (44.85 mg/kg) was recorded from Damu Degele kebele, followed by Warabu Masi (44.16 mg/kg). In comparison, a lesser amount of HMF (32.55 mg/kg) was obtained from the Fitte Wato kebele collected sample with an overall mean of 40.52 mg/kg. The presence of elevated levels of HMF in honey collected from Damu Degele and Worabu Messe suggests that the honey has undergone aging or extended storage in its natural containers. It is important to note that the honey samples used in the present study were freshly harvested and processed, indicating that the observations made by Shapla *et al*. (2018) are not applicable to the current research.

The present study revealed a WISC (estimation for the presence of impurities in honey) ranging from 0.24 to 0.87% with an overall mean value of 0.59%. There was a statistically significant difference among SBH, which was collected from the three kebeles. It was observed that considerable amounts of WISC (0.87%) had been obtained for the SBH sample, which was collected from Damu Degele, followed by Worabu Messe (0.67%), while a lesser amount of WISC (0.24%) was obtained from the SBH sample, which was collected from the Fite Wacho site. Our result showed that the honey samples collected from Fite Wacho were the cleanest of all other samples. Besides, the overall mean value of this result is slightly lower (0.59%) than the mean value (0.69%) reported in the previous study (Gela *et al.,* 2021).

### 3.2. Sugar profile of SBH

In the present investigation, an analysis of sugars in stingless bee honey sourced from the Wonchi district of South West Shewa, Ethiopia, revealed the presence of three distinct types: fructose, glucose, and sucrose (**Figure 2**). Among these, fructose emerged as the predominant sugar, constituting the highest proportion at 18.87 g per 100 g (53.87%), followed by glucose (11.10 g/100 g), while sucrose exhibited the lowest concentration, accounting for 5 g per 100 g (14.33%). Furthermore, the glucose content in our current study, standing at 11.10 g/100 g, is observed to be lower than the outcomes reported by Teferi *et al*. (2022), who documented a value of 27.67 g/100 g. This suggests a distinct variation in the carbohydrate composition of the studied samples in comparison to previous research conducted in Ethiopia and the Amazon Region of Brazil.

**Figure 2:**
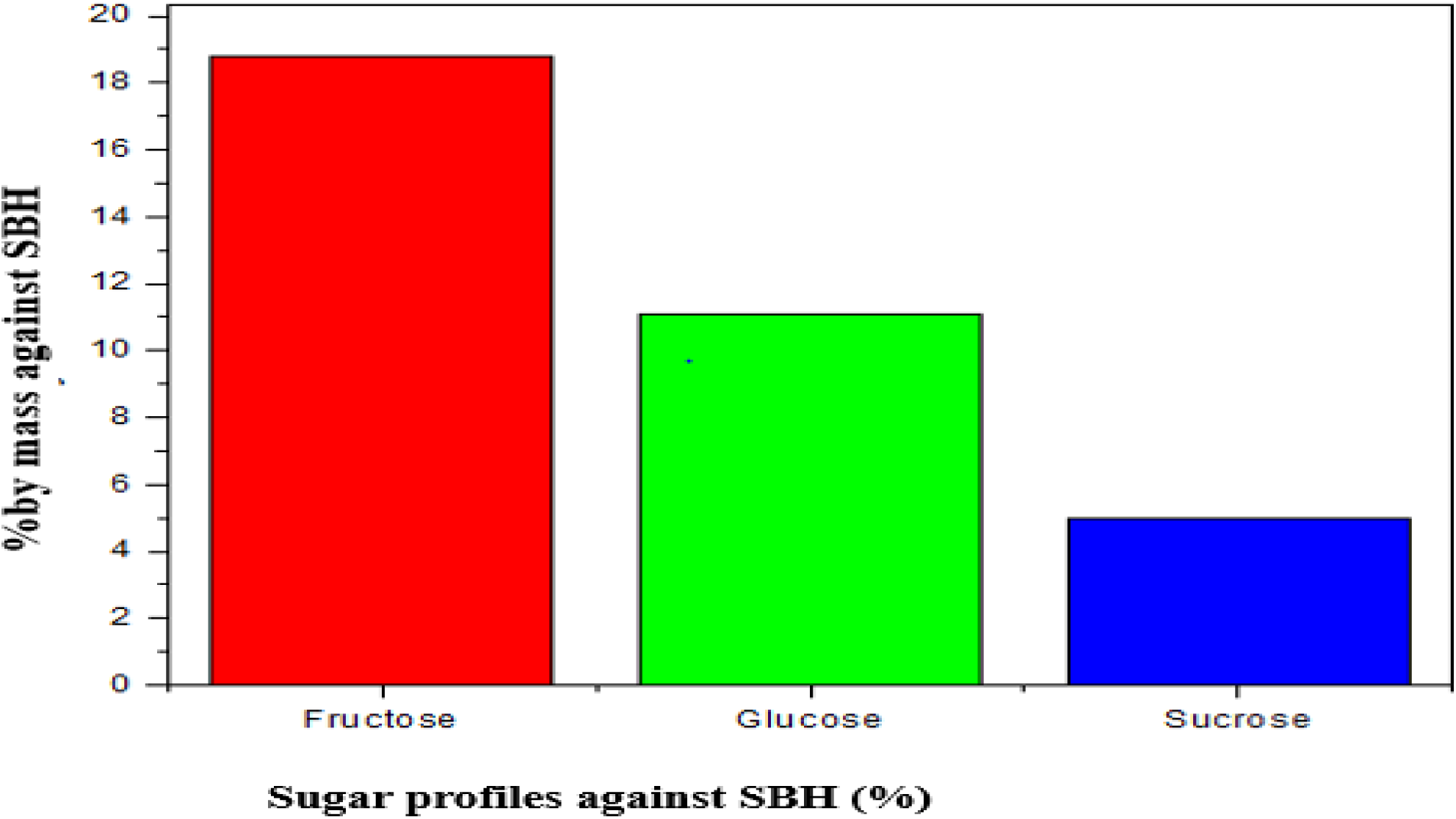
The sugar profile of SBH collected from Wonchi District in 2023

Several investigations have proposed that the sugar composition in honeybees can be influenced by geographic locations, impacting the nectar sources from various plants, as well as environmental factors like humidity, temperature, postharvest honey processing, and storage duration (Biluca *et al.,* 2016). This finding supports the notion that in high-quality stingless bee honey, fructose tends to be more prevalent compared to glucose and sucrose.

### 3.3. Determination of Mineral Content

Variability in mineral composition was observed across different honey sites, with K being the most prevalent mineral. Concentrations ranged from 14.5b ± 0.41 mg/kg to 18.34a ± 0.30 mg/kg (**Table 2**). The highest K content, recorded at 19.21 ± 1.84 mg/kg, was found in honey from the Damu Dagele kebele. In contrast, Mg exhibited the lowest content, ranging from 4.540 ± 0.867 mg/kg, sourced from the Fite Wacho kebele. Ca ranked as the second-highest mineral, with a measured value of 8.234 ± 1.656 mg/kg, obtained from the Warabu Messee, Wonchi District. The mineral contents of these minerals varied substantially among the study locations. The honey mineral content is affected by flora geographical and botanical origin, bees type and activity, extraction technique and storage conditions (Nnadi, 2020). Our findings here are quite similar to those of (Gela *et al*. 2021), who reported ash contents ranging from 0.21% to 0.57% for stingless bee honey. This is important because the mineral content in honey has far-reaching effects on many things.

**Table 2:** Determination of mineral content in SBH samples.

| Sampling locations | Mean ± SD | | |
| --- | --- | --- | --- |
|  | Mg | K | Ca |
| Damu Degele | $4.37^a \pm 0.26$ | $18.34^{a\pm} 0.30$ | $6.84^{a\pm} 0.37$ |
| Fite Wacho | 2.467<sup>c</sup> ±0.19 | 14.5<sup>b</sup>±0.41 | 4.16<sup>b</sup>±0.47 |
| Worabu Messe | 3.634<sup>b</sup> ±0.06 | 17.09<sup>a</sup>±0.06 | 7.40<sup>a</sup>±0.01 |
| Total | 3.48±0.17 | 16.64±0.257 | 6.13±0.283 |
| LSD | 0.52 | 1.67 | 1.61 |
| CV% | 7.54 | 5.03 | 13.2 |
| P = value | 0 | 0 | 0 |

*CV=Coefficient of variation, SD=Standard Deviation, LSD=Least Significant Difference; and superscripts in the column followed by the different letter are significantly different at the 5% level by Fisher's LSD.*

The use of this agent as marker can help to determine the geographical source of honey, allowing for their tracing and authentication (*Zhang et al. 2023*). This also correlates with the prevalent floral sources exploited by bees that are known to sequester different mineral compositions in plant nectar (Machado *et al*. 2018). Additionally, the mineral composition can be used to evaluate internal quality and authenticity measures of honey due to their natural profile may vary as a result of adulteration or improper processing (Downey, 2003). Furthermore, the ash content of stingless bee honey also differs geographically, with Southeast Asia reporting values between 0.1% and 0.3% (Tey *et al*. 2022), and South America finding it to be 0.2% to 0.5% (Cavalcante *et al*. 2020).

### 3.4. Correlation Between Mineral Content

The analysis of mineral content correlations in SBH reveals a notably strong and statistically significant relationship between Ca and Mg, denoted by a perfect significance level (**Table 3**). Additionally, a substantial correlation was observed between K and Mg, with a correlation coefficient of 0.72. Similarly, the correlation between K and Ca was highly significant, exhibiting a correlation coefficient of 0.65.

**Table 3.** Correlation between mineral content.

| Variables | Mg | Ca | K |
| --- | --- | --- | --- |
| Mg |  |  |  |
| Ca | 0.88 |  |  |
| K | 0.73 | 0.66 |  |

*Correlation is significant at the 0.05 level*

### 3.5. Correlation Between Physicochemical and Proximate Analysis

The correlation analysis result for all physicochemical and proximate analysis were the highest correlation except fatty and Ash (**Table 4**). The correlation between fatty with other physicochemical and proximate analyses showed very insignificant correlation such as fatty, total water soluble, fatty with pH, fatty with electrical conductivity, fatty with moisture and fatty with water-soluble solid content. The other insignificant correlation observed on ash mineral content with other physicochemical and proximate analyses, such as ash with free acidity and ash with protein.

**Table 4.** Correlation between physicochemical and proximate analysis of SBH.

| Variables | TSS | PH | HMF | EC | acidity | WISC | Moisture<br>contents | fatty | protein | Ash |
| --- | --- | --- | --- | --- | --- | --- | --- | --- | --- | --- |
| TSS |  |  |  |  |  |  |  |  |  |  |
| PH | 0.94* |  |  |  |  |  |  |  |  |  |
| HMF | 0.78* | 0.73* |  |  |  |  |  |  |  |  |
| EC | 0.93* | 0.87* | 0.81* |  |  |  |  |  |  |  |
| Free acidity | 0.81* | 0.66* | 0.56* | 0.7* |  |  |  |  |  |  |
| Water insolubility | 0.89* | 0.85* | 0.86* | 0.89* | 0.55* |  |  |  |  |  |
| Moisture | 0.85* | 0.90* | 0.69* | 0.88* | 0.63* | 0.80* |  |  |  |  |
| Fatty | 0.48 | 0.41 | 0.52 | 0.37 | 0.77* | 0.22 | 0.31 |  |  |  |
| Protein | 0.76* | 0.66* | 0.53* | 0.62* | 0.90* | 0.57* | 0.68* | 0.7* |  |  |
| Ash | 0.64* | 0.74* | 0.55* | 0.67* | 0.45 | 0.54* | 0.90* | 0.28 | 0.50* |  |

*EC=Electric conductivity; HMF=Hydroxyl-Methyl Furfural; WISC=Water-Insoluble Solids Content; and \** *indicates $p < 0.05$ (correlation is significant).*

## 4. CONCLUSION AND RECOMMENDATIONS

The present study assessed the physicochemical properties and proximate analysis of stingless bees’ honey from the southwest Shewa Zone, Ethiopia. The stingless bee honey samples analyzed in this study exhibited significant richness in carbohydrates, proteins, and minerals. The identification of such molecules in SBH suggests that SHB in the study area has high nutritional quality. The physicochemical attributes of honey from nearly all locations fell within the prescribed limits set by Ethiopian, Codex, and EU standards. The study highlights the physicochemical profile of SBH, which is characterized by significantly higher levels of acidity (8.62±2.73 meq/kg), pH (3.39±0.32), electrical conductivity (1.8±0.52), moisture content (20.12±1.14 g/100 g), and hydroxymethylfurfural (HMF) content (40.52±6.61). These attributes collectively indicate the quality and freshness of SBH. Low pH enhances the shelf life of SBH by inhibiting the growth and presence of microorganisms, and it may also provide potential medicinal benefits. The acidity, which serves as a quality parameter indicating the presence of organic acids in honey, exhibited variability among the three kebeles included in this study. The study’s findings also indicate the area’s potential for producing quality honey and highlight the ease of identifying priority areas for stingless bee conservation. Based on the findings of the present study, it is recommended that a modern honey production system be implemented to improve honey quality, market access, production efficiency, and overall sustainability, while beekeepers should receive focused conservation training programs to ensure the long-term survival of stingless bees and their valuable products.

## Acknowledgment

We extend our gratitude to Wolkite University for their invaluable support for this research paper.

## Authors’ contributions

**S.A:** Conceptualization, methodology, writing original-draft, and validation; **D.L:** Writing-review/research & editing, visualization, resources, data curation, and facilitating; investigation, editing, and supervision.

